# Impact of cobas PCR Media Freezing on SARS-CoV-2 Viral RNA Integrity and Whole Genome Sequencing Analyses

**DOI:** 10.1101/2021.02.05.430022

**Authors:** Patrick Benoit, Floriane Point, Simon Gagnon, Daniel E. Kaufmann, Cécile Tremblay, Richard Paul Harrigan, Isabelle Hardy, François Coutlée, Simon Grandjean Lapierre

## Abstract

SARS-CoV-2 whole genome sequencing is an important molecular biology tool performed to support many aspects of the response to the pandemic. Freezing of primary clinical nasopharyngeal swab samples and shipment to reference laboratories is usually required since RNA sequencing is rarely available in routine clinical microbiology laboratories where initial diagnosis and support to outbreak investigations occur. The cobas PCR Media transport medium developed by Roche facilitates high throughput analyses on cobas multianalyzer PCR platforms. There is no data on the stability of SARS-CoV-2 RNA after freezing and thawing of clinical samples in this transport medium, but potential denaturing of the molecular template could impair test results. Our objective was to compare the quality and results of SARS-CoV-2 genomic sequencing when performed on fresh or frozen samples in cobas PCR Media. Viral whole genome sequencing was performed using Oxford Nanopore Technologies MinION platform. Genomic coverage and sequencing depth did not significantly differ between fresh and frozen samples (n=10). For samples with lower viral inoculum and PCR cycle threshold above 30, sequencing quality scores and detection of single nucleotide polymorphisms did not differ either. Freezing of cobas PCR Media does not negatively affect the quality of SARS-CoV-2 RNA sequencing results and it is therefore a suitable transport medium for outsourcing sequencing analyses to reference laboratories. Those results support secondary use of diagnostic nasopharyngeal swab material for viral sequencing without requirement for additional clinical samples.

## INTRODUCTION

SARS-CoV-2 viral genomic sequencing plays an important role in the short and long-term responses to the COVID-19 pandemic including global and national surveillance of the virus evolution, understanding of SARS-CoV-2 natural history of disease and outbreak investigations (1-4). Viral whole genome sequencing primarily occurs in reference laboratories and is rarely performed where clinical diagnosis or outbreak investigations happen. Therefore, freezing of primary samples is required prior to viral genomic amplification and sequencing.

cobas PCR Media is a transport medium developed by Roche that simplifies linkage between pre-analytical sampling and analytical testing and is adapted for high throughput analyses on cobas multianalyzer PCR platforms. It contains guanidine hydrochloride which is a denaturing agent used to dissociate nucleoproteins and inactivate RNases. The manufacturer does not recommend freezing the cobas PCR Media because of risks of molecular template denaturation (5). Freezing of other transport media was previously shown not to negatively impact the detection of SARS-CoV-2 by RT-PCR (6). However, it is unknown whether, and how, freezing of cobas PCR Media indeed denatures SARS-CoV-2 RNA and if it negatively affects viral genomic sequencing.

In this study, we compared the quality and results of SARS-CoV-2 whole genome sequencing between fresh and frozen samples obtained from combined oral and nasopharyngeal swabs (ONPS). We used matched split samples collected in cobas PCR Media and either processed following collection and storage at 4 °C or frozen for one week at −80 °C and thawed prior to sequencing. Our protocol and analysis address the necessity for most clinical microbiology laboratories to refer frozen primary clinical samples that were used for diagnostic purposes to reference laboratories in order to access viral genomic information.

## MATERIALS AND METHODS

### CLINICAL SAMPLES

Ten clinical samples were included in this study. All samples were a combined ONPS submitted to our laboratory in cobas PCR Media initially found to be positive for SARS-CoV-2 using the FDA emergency use authorization (EUA) approved and validated cobas 8800 automated RT-PCR system which simultaneously tests both the ORF1 a/b and E-gene viral molecular targets (7). To assess the impact of viral load or initial amount of RNA template on sequencing, we purposively included samples testing positive at a broad range of cycle threshold (Ct) from 14.4 to 34.7 for the ORF1 a/b target. This strategy ensured inclusion of samples with low (high Ct) and high (low Ct) viral loads.

### FREEZING AND THAWING

Following initial positive RT-PCR testing, samples were split in equal volumes. The first half of each samples was maintained at 4 °C, according to manufacturer’s recommendation, and then processed for whole genome sequencing as described below. The second half of each sample was frozen at −80 °C for 7 days and thawed. RT-PCR was then repeated using the same cobas 8800 system and viral genomic sequencing was performed using identical methods.

### SARS-CoV-2 GENOMIC SEQUENCING

Viral RNA was extracted from 0.2 mL of cobas PCR Media using Maxwell® 16 instrument (Promega, Madison, WI, USA) for final elution in 30μL. Viral whole genome sequencing was performed using the ARTIC Network V3 protocol on Oxford Nanopore Technologies (ONT) (Oxford, United Kingdom) MinION® long read sequencing platform. Since its initial publication online in January 2020, the ARTIC protocol has become one of the most widely used approach to SARS-CoV-2 genomic sequencing. This protocol has yielded a significant sequence contribution to the GISAID global database and is currently used for surveillance by many public health agencies (8, 9). Briefly, genome amplification was performed by reverse transcriptase multiplex PCR using nCoV-2019 V3 primer combinations (Integrated DNA Technologies). This set of primers was previously shown to produce high genomic coverage with low variance on the whole viral genome (8). RT-PCR amplicons were assessed by Qubit® fluorometric DNA Quantification (Thermo Fisher Scientific, Waltham, MA, USA). For samples with post RT-PCR DNA quantity below 250 ng, we omitted the dilution step of the sample in 45 μL of molecular grade water before library preparation as recommended in the ARTIC protocol. Such low inoculums were observed in three samples with both ORF1 a/b and E-gene targets Cts over 30 (samples 1, 2, 3). Sequencing libraries were prepared following ONT protocol for genomic DNA with native barcoding and using 9.4.1 flow cells on the MinION® platform. Raw sequencing reads fast5 files were base called with high accuracy using ONT proprietary software Guppy (v3.4.5). Reads were demultiplexed and filtered using the online available ARTIC network bioinformatic pipeline solution (10). This filtering process includes exclusion of sequencing reads respectively below 400 and above 700 base pairs which do not correspond to expected amplicons length resulting from the RT-PCR primer set. Reads were mapped to the Wuhan-Hu-1 SARS-CoV-2 reference genome (GeneBank accession number MN908947.3) using minimap2 (v2.17). Predominantly sequenced nucleotides at positions for which a minimal depth of 20 reads had been achieved were used to generate consensus viral genomic sequences. Potential subpopulations or mixed infections were not considered, and hence a unique consensus sequence was generated for each isolate.

## DATA ANALYSIS

We compared mean sequencing Q-scores with corresponding error rates and accuracy, single nucleotide polymorphisms (SNPs) identification and diversity of sequenced alleles on identified SNP genomic positions. We used those later metrics as surrogate markers of post-freezing viral RNA integrity. Q-scores represent ONT’s sequencing platform and base calling software internal assessment of sequencing read quality. The Q-score of a given base is defined as Q = 10log_10_(e) where (e) is the estimated probability of the base call being wrong. We used a two-tailed paired samples t-test with an alpha value of 0.05 to compare pre- and post-freezing variables. All statistical analyses were performed using GraphPad Prism (San Diego, CA USA).

To simulate prospective outbreak investigation, we supplemented the pre- and post-freezing sequence datasets with a back catalog of 50 SARS-CoV-2 genomic sequences from our institution (unpublished data) hence generating two mocked nosocomial viral pangenomes. We independently analyzed both augmented data sets as if searching for potential transmission clusters. Consensus sequences were compared, and phylogenetic trees were built using UGENE (v37) with the PHYLIP Neighbor Joining method without bootstrapping. To simulate national surveillance and assessment of circulating viral clades, we independently compared the pre- and post-freezing sequence datasets with published and well described SARS-CoV-2 reference genomes submitted to Nextstrain (https://nextstrain.org/sars-cov-2/) (11).

All laboratory testing including sequencing and data analyses were performed in Centre Hospitalier de l’Université de Montréal. Patients’ symptoms nature and relative temporality with clinical sampling, or potential person to person transmission events were not taken into consideration.

## DATA AVAILABILITY

SARS-CoV-2 sequences from this study are available at GenBank under continuous accession numbers MW309425 to MW309442.

## RESULTS

### RT-PCR

Upon initial testing after maintenance of clinical samples at 4 °C in cobas PCR Media, RT-PCR Cts ranged from 14.4 to 34.7 and 14.9 to 34.9 respectively for the ORF1 a/b and E-gene targets. After freezing for 7 days at −80 °C, RT-PCR Cts ranged between 17.8 to 31.8 and 17.9 to 33.8 for the same targets. No statistically significant difference was observed between pre- and post-freezing Cts for the ORF1 a/b target (*p-value* 0.64). One sample only became positive on the E-gene target after freezing. Excluding this sample from the analysis, post-freezing Cts for the E-gene target were 1.1 Ct higher after freezing (*p-value* 0.01) (Table 1).

**Table 1 :** Impact of cobas PCR Media freezing on SARS-CoV-2 RT-PCR Ct levels. Difference in SARS-CoV-2 RT-PCR Ct levels after 7-day freezing in cobas PCR Media. Samples are presented in decreasing order of Ct value on pre-freezing ORF1 a/b RT-PCR.

|  | **PRE** |  | **POST** |  | **DELTA (PRE - POST)** |  |
| --- | --- | --- | --- | --- | --- | --- |
|  | ORF1 a/b | E-gene | ORF1 a/b | E-gene | ORF1 a/b | E-gene |
| | CT | CT | CT | CT | $\Delta$ CT | $\Delta$ CT |
| 1 | 34.7 | NEG | 31.8 | 33.8 | 3.0 | N/A |
| 2 | 32.2 | 34.9 | 30.4 | 34.5 | 1.8 | 0.4 |
| 3 | 30.9 | 32.4 | 30.8 | 33.2 | 0.1 | -0.8 |
| 4 | 28.5 | 29.4 | 28.3 | 29.4 | 0.2 | -0.1 |
| 5 | 26.1 | 26.5 | 26.4 | 27.4 | -0.3 | -0.9 |
| 6 | 23.2 | 23.3 | 24.0 | 24.5 | -0.8 | -1.2 |
| 7 | 21.3 | 21.6 | 23.0 | 23.2 | -1.7 | -1.6 |
| 8 | 18.9 | 19.2 | 20.4 | 20.9 | -1.6 | -1.8 |
| 9 | 16.7 | 16.6 | 16.9 | 17.4 | -0.1 | -0.8 |
| 10 | 14.4 | 14.9 | 17.8 | 17.9 | -3.4 | -3.0 |
| Mean | 24.7 | 24.3 | 25.0 | 26.2 | -0.3 | -1.1 |
| *p-value* |  |  |  |  | 0.64 | 0.01 |

### VIRAL GENOMIC SEQUENCING

No statistically significant difference was observed between the sequencing yields before or after freezing. Indeed, freezing did not negatively impact the total number of sequenced bases and mapped reads with pre- / post-freezing mean deltas of 11 Mb (*p-value* 0.57) and 938 reads (*p-value* 0.31) for those key metrics. Also importantly, 20X sequencing depth, allowing for wild type or variant allelic identification within our protocol, was achieved for an average of 83.9% and 83.7% of the viral genome respectively before and after freezing (*p-value* 0.90) (Table 2). Such similarity was also observed for all other evaluated depth thresholds (1X, 5X, 10X, 50X). As expected, sequencing data yield, depth and coverage were inversely correlated to the Ct value both in pre- (*p-value* 0.0007) and post-freezing (*p-value* 0.0003) samples. Less sequencing data was hence generated in the sub-group of low viral inoculum and high Ct samples 1, 2 and 3 but freezing did not negatively impact sequencing yields in this subgroup either (Fig. 1).

**Table 2.** Impact of freezing on SARS-CoV-2 Genomic Sequencing Data Yield. Difference in SARS-CoV-2 sequencing generated bases, reads and corresponding genomic coverage at various depth thresholds after 7-day freezing in cobas PCR Media. Samples are presented in decreasing order of Ct value on pre-freezing ORF1 a/b RT-PCR.

|  | PRE |  |  |  | POST |  |  |  | DELTA (PRE-POST) |  |  |  |
| --- | --- | --- | --- | --- | --- | --- | --- | --- | --- | --- | --- | --- |
|  | Bases | Mapped reads | 20X | Average depth | Bases | Mapped reads | 20X | Average depth | Bases | Reads | 20X | Average depth |
|  | Mb | n | % | n | Mb | n | % | n | Mb | n | Δ % | Δ n |
| 1 | 16.7 | 151 | 2.6 | 2 | 17.9 | 638 | 12.6 | 8 | -1.2 | -487 | -10.0 | -6 |
| 2 | 19.1 | 5295 | 55.9 | 66 | 14.8 | 4950 | 51.4 | 63 | 4.3 | 345 | 4.5 | 4 |
| 3 | 39.4 | 18782 | 87.3 | 236 | 19.9 | 10858 | 81.6 | 136 | 19.5 | 7924 | 5.7 | 100 |
| 4 | 72.6 | 32491 | 98.4 | 414 | 80.9 | 33389 | 98.0 | 426 | -8.3 | -898 | 0.4 | -12 |
| 5 | 68.4 | 34382 | 97.7 | 440 | 125.7 | 36137 | 98.6 | 461 | -57.3 | -1755 | -0.8 | -21 |
| 6 | 179.4 | 38324 | 99.2 | 489 | 115.8 | 36751 | 99.2 | 470 | 63.6 | 1573 | 0.0 | 19 |
| 7 | 140.7 | 38000 | 98.6 | 483 | 176.9 | 38155 | 98.6 | 483 | -36.2 | -155 | 0.0 | 0 |
| 8 | 216.2 | 37508 | 99.3 | 477 | 142.3 | 37133 | 98.5 | 474 | 73.9 | 375 | 0.8 | 3 |
| 9 | 140.7 | 38534 | 100.0 | 492 | 204.2 | 38626 | 100.0 | 494 | -63.5 | -92 | 0.0 | -2 |
| 10 | 217.4 | 39628 | 100.0 | 499 | 102.7 | 37075 | 99.0 | 471 | 114.7 | 2553 | 1.0 | 27 |
| Mean | 111.1 | 28310 | 83.9 | 360 | 100.1 | 27371 | 83.7 | 348 | 11.0 | 938 | 0.2 | 11 |
| p-value |  |  |  |  |  |  |  |  | 0.57 | 0.31 | 0.90 | 0.33 |

**Figure 1.**
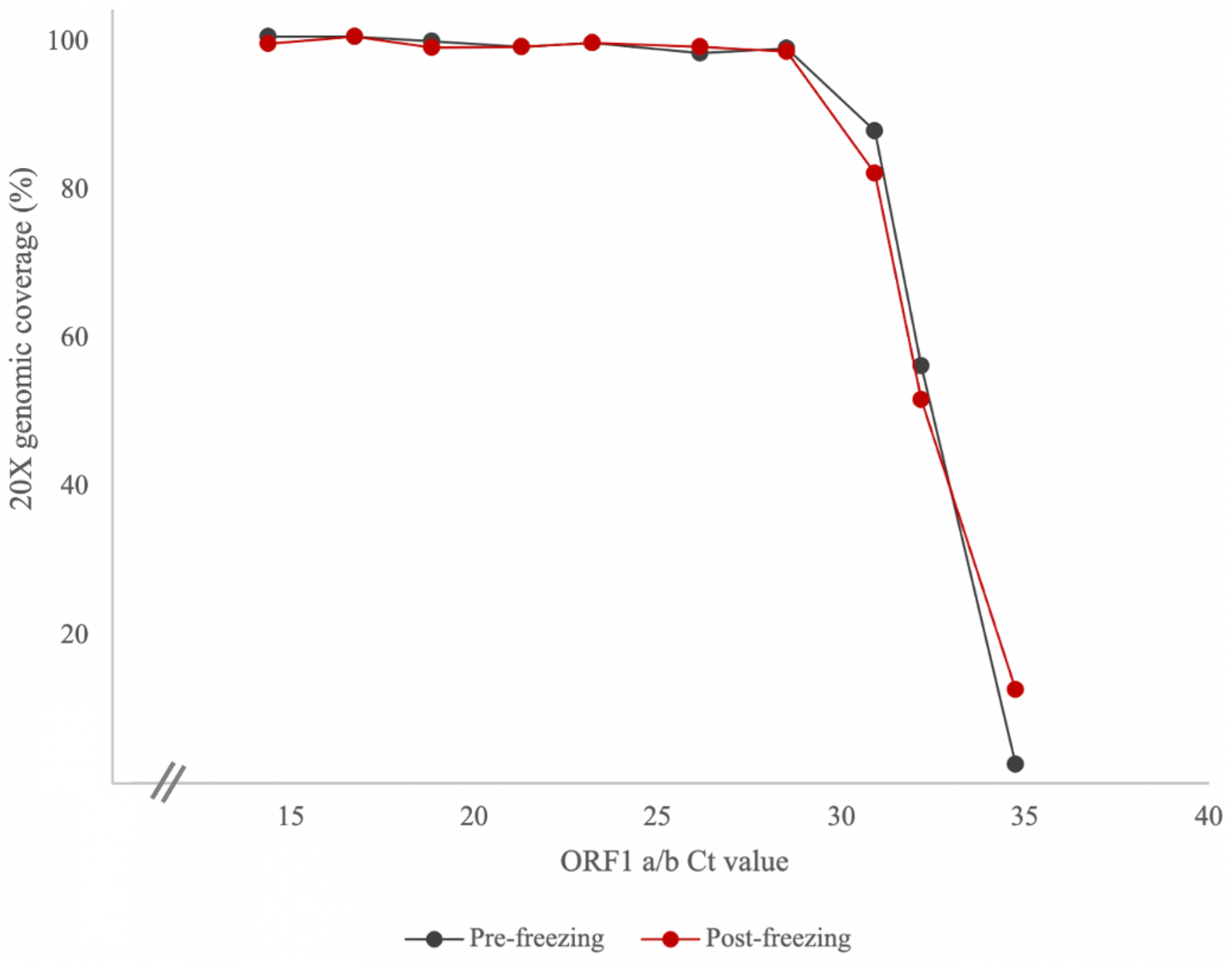
ORF1 a/b Ct value correlation with 20X genomic sequencing coverage. Coverage of genome at 20X in relation to pre-freezing Ct value for ORF1 a/b. Less sequencing data was generated with samples with higher cycle thresholds (Ct) but freezing did not negatively impact sequencing yields.

No statistically significant decrease was observed in Q-scores (*p-value* 0.07) and base call accuracy (*p-value* 0.10) after freezing (Table 3). Except for samples 1 (ORF1 a/b Ct 34.74) and 2 (ORF1 a/b Ct 32.16), freezing did not impact SNP detection and identified mutations were identical in both sequencing analyses. Looking in more depth at each single read for those specific mutation sites, the percentage of alternate bases leading to SNP calling did not significantly change after freezing (*p-value* 0.31). This ratio of variant versus wild type alleles at each mutation site was the same before and after freezing and suggests that the viral molecular template was not significantly degraded by the freezing process in cobas PCR Media.

**Table 3.** Impact of cobas PCR Media freezing on SARS-CoV-2 Genomic Sequencing Accuracy. Differences in SARS-CoV-2 sequencing accuracy and single nucleotide polymorphisms identification after 7-day freezing in cobas PCR Media. Samples are presented in decreasing order of Ct value on pre-freezing ORF1 a/b RT-PCR.

|  | PRE |  |  |  |  | POST |  |  |  |  | DELTA (PRE - POST) |  |  |  |  |
| --- | --- | --- | --- | --- | --- | --- | --- | --- | --- | --- | --- | --- | --- | --- | --- |
|  | Q-Score | Accuracy | Non-calls | SNP | Alternate allele | Q-Score | Accuracy | Non-calls | SNP | Alternate allele | Q-Score | Accuracy | Non-calls | SNP | Alternate allele |
|  |  | % | n | n | % | N/A | % | n | n | % |  | Δ % | Δ n | Δ n | Δ % |
| 1 | 12.9 | 99.2 | 27306 | 0 | N/A | 13.0 | 99.3 | 21402 | 3 | 89.1 | -0.1 | 0.0 | 5904 | -3 | N/A |
| 2 | 13.1 | 99.3 | 4379 | 9 | 72.2 | 12.8 | 99.1 | 2570 | 6 | 86.1 | 0.3 | 0.2 | 1809 | 3 | -13.9 |
| 3 | 13.3 | 99.4 | 664 | 18 | 84.7 | 12.9 | 99.2 | 1011 | 18 | 84.4 | 0.4 | 0.2 | -347 | 0 | 0.3 |
| 4 | 13.7 | 99.6 | 251 | 15 | 84.1 | 13.7 | 99.6 | 12 | 15 | 87.1 | 0.0 | 0.0 | 239 | 0 | -2.9 |
| 5 | 13.7 | 99.6 | 3 | 16 | 89.8 | 13.6 | 99.5 | 1 | 16 | 89.2 | 0.1 | 0.0 | 2 | 0 | 0.7 |
| 6 | 13.6 | 99.5 | 0 | 11 | 88.0 | 13.4 | 99.5 | 0 | 11 | 87.8 | 0.2 | 0.1 | 0 | 0 | 0.2 |
| 7 | 13.6 | 99.6 | 0 | 16 | 89.4 | 13.6 | 99.5 | 0 | 16 | 88.8 | 0.0 | 0.0 | 0 | 0 | 0.5 |
| 8 | 13.6 | 99.5 | 0 | 21 | 88.9 | 13.6 | 99.5 | 0 | 21 | 88.2 | 0.0 | 0.0 | 0 | 0 | 0.8 |
| 9 | 13.7 | 99.6 | 0 | 21 | 89.3 | 13.6 | 99.5 | 0 | 21 | 89.4 | 0.0 | 0.0 | 0 | 0 | -0.2 |
| 10 | 13.6 | 99.5 | 0 | 14 | 89.3 | 13.5 | 99.5 | 0 | 14 | 90.2 | 0.1 | 0.0 | 0 | 0 | -0.9 |
| Mean | 13.5 | 99.5 |  | 14 | 86.2 | 13.4 | 99.4 |  | 14 | 88.0 | 0.1 | 0.0 | 761 | 0 | -1.7 |
| p-value |  |  |  |  |  |  |  |  |  |  | 0.07 | 0.10 | 0.24 | 1.00 | 0.31 |

In the mocked outbreak investigation, samples with higher genomic similarity were identified. Although our study was not a molecular epidemiology study and did not include clinical correlation with putative transmission events, those molecular clusters were identical in both pre- and post-freezing analyses (Fig. 2). For the surveillance clade typing simulated application, phylogenetic placement was also identical in the pre- and post-freezing mocked data sets (Fig. 3). For viral clade typing and comparison to reference genomes, samples 1 and 2 could not be included in the analysis because of too small genomic coverage. All phylogenetic placement results were expected and in agreement with previously described findings on identical SNP typing.

**Figure 2.**
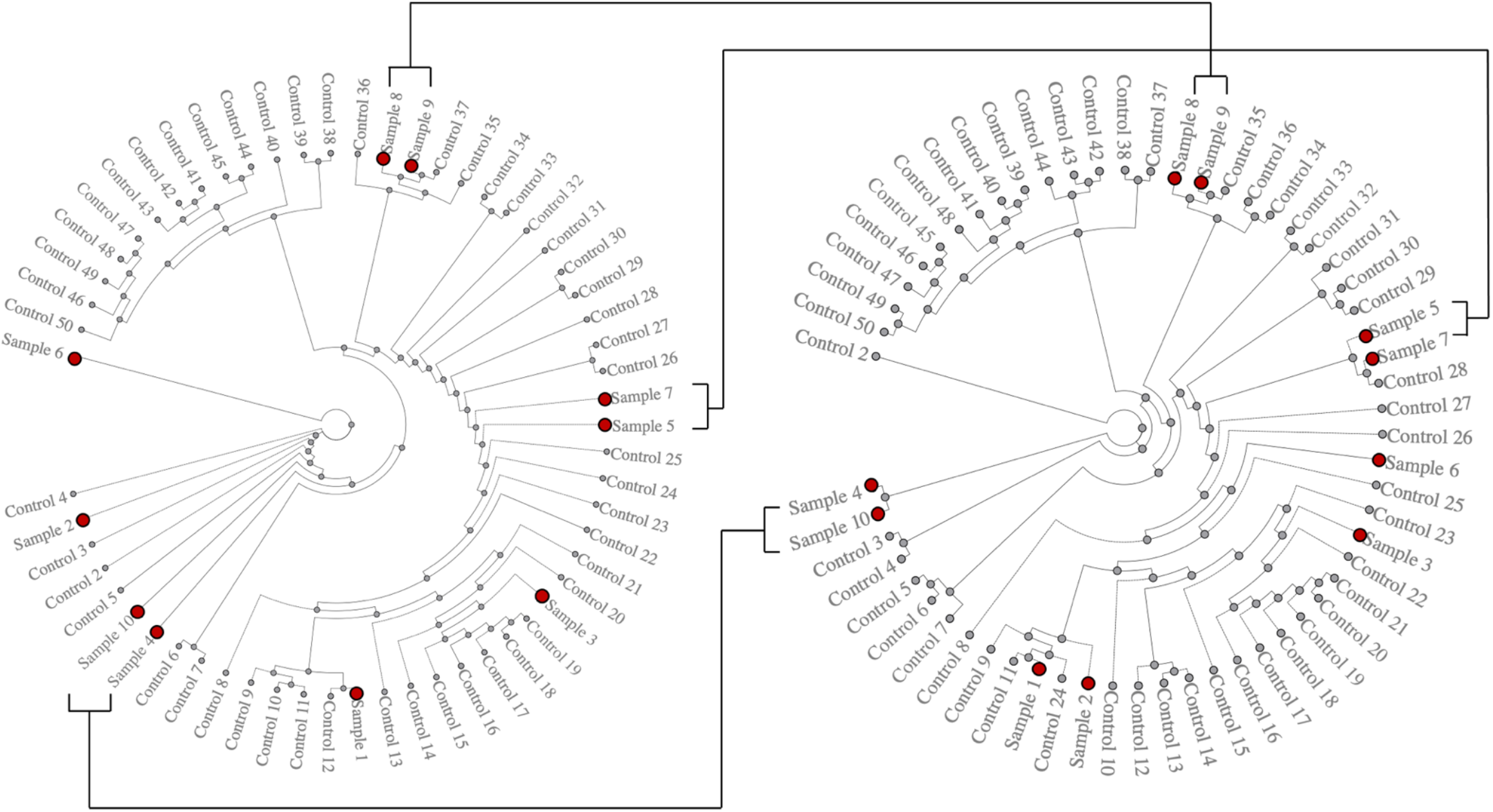
Simulated SARS-CoV-2 outbreak investigation. A simulated outbreak investigations with a back catalog of 50 SARS-CoV-2 genomic sequences from our institution (unpublished data). Phylogenetic trees were constructed using UGENE (v37) with the PHYLIP Neighbor Joining method without bootstrapping. Pre- (left) and post-freezing (right) genomic sequences show identical potential outbreak clusters within our samples.

**Figure 3.**
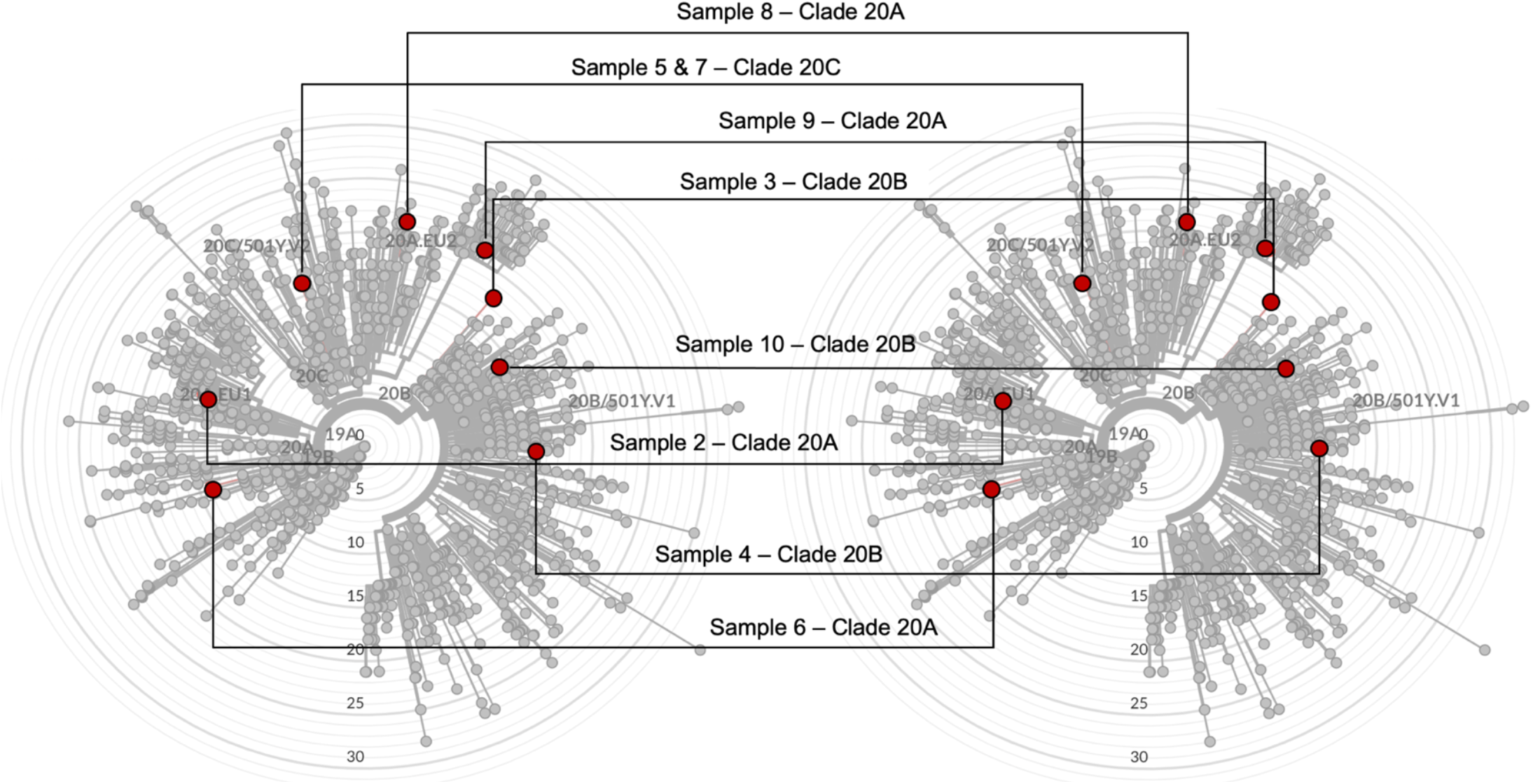
SARS-CoV-2 Study Isolates Placement Within Global Reference Sequences. Radial rooted phylogenetic representation of SARS-CoV-2 reference sequence submitted to Nextstrain (https://nextstrain.org/sars-cov-2/). Pre- (left) and post-freezing (right) genomic sequences (red) showing identical phylogenetic placement within global reference sequences (grey).

## DISCUSSION

In our study, a single freeze - thaw cycle of ONPS clinical samples in cobas PCR Media did not significantly impact analytical sensitivity of SARS-CoV-2 RT-PCR on cobas 8800® automated system on this limited set of samples. Although we observed a Ct increase of 1.1 (*p-value* 0.01) after freezing for the E-gene target, one of our samples was also found to be positive for this target only after freezing. Freezing the sample in cobas PCR Media did not degrade viral molecular templates and did not negatively affect viral genomic sequencing analyses. Khiri *et al* had previously shown that freezing of cervical samples in cobas PCR Media did not negatively impact the sensitivity of PCR for human papillomavirus detection (12). Our study confirms this holds true for SARS-CoV-2. To our knowledge, this is the first study to formally evaluate the impact of freezing clinical samples in cobas PCR Media for downstream sequencing analyses either for SARS-CoV-2 or for any other target pathogen or molecular template. Our study confirms the ability of cobas PCR Media to maintain SARS-CoV-2 genomic RNA at −80 °C for subsequent sequencing analyses. Note that the PCR amplicons generated in this study are relatively small (∼400 bp), so this protocol may be more robust to RNA damage than methods which require long, intact starting molecules. Our results should also not be generalized to other transport media without independent confirmation.

Our study included three samples with RT-PCR Cts above 30.0 which are considered to have a lower viral load. For those samples, SNP calling showed variability and genomic coverage was insufficient to allow detailed phylogenetic analyses. This phenomenon was observed both before and after freezing and is hence believed to be due to low viral inoculum rather than transport medium related viral RNA denaturation. Our study included only 10 samples but the extensive comparability between pre- and post-freezing sequencing results suggests that a higher denominator would not have led to different conclusions. It is possible that a freezing period longer than 7 days would have led to worse sequencing results after thawing but our protocol did not assess such longer-term effect. Seven days represents a sufficient delay for transportation to reference laboratories performing viral sequencing and our study hence provides meaningful information to clinical laboratories involved in routine diagnostic testing.

## CONCLUSION

Our study demonstrates that the freezing of cobas PCR Media at −80 °C does not affect viral genomic sequencing quality and results for SARS-CoV-2. The consistent results between pre- and post-freezing support potential secondary use of diagnostic oral and nasopharyngeal swab material for viral sequencing without requirement for additional clinical sampling. Our findings will simplify the collection and storage of samples in laboratories where this transport medium is utilized.

## ACKNOWLEDGEMENT

This study was funded by the *Réseau SIDA-Maladies Infectieuses* of the *Fond de Recherche Santé Québec*, Roche Diagnostics (Laval, Canada) and CIHR/CITF grant VR2-173203. The funders had no role in study design, data collection and interpretation, or the decision to submit the work for publication.

